# Nitric oxide synthase neurons in the preoptic hypothalamus are sleep-active and contribute to regulating NREM and REM sleep and lowering body temperature

**DOI:** 10.1101/2021.05.14.444161

**Authors:** Edward C. Harding, Wei Ba, Reesha Zahir, Xiao Yu, Raquel Yustos, Bryan Hsieh, Leda Lignos, Alexei L. Vyssotski, Timothy Constandinou, Nicholas P. Franks, William Wisden

## Abstract

When mice are exposed to external warmth, nitric oxide synthase (NOS1) neurons in the median and medial preoptic (MnPO/MPO) hypothalamus induce sleep and concomitant body cooling. However, how these neurons regulate baseline sleep and body temperature is unknown. Using calcium photometry, we show that NOS1 neurons in MnPO/MPO are predominantly NREM active. This is the first instance of a predominantly NREM-active population in the PO area, or to our knowledge, elsewhere in the brain. In addition to releasing nitric oxide, NOS1 neurons in MnPO/MPO can release GABA, glutamate and peptides. We expressed tetanus-toxin light-chain in MnPO/MPO NOS1 cells to reduce vesicular release of transmitters. This induced changes in sleep structure: over 24 hours, mice had less NREM sleep in their dark (active) phase, and more NREM sleep in their light (sleep) phase. REM sleep episodes in the dark phase were longer, and there were fewer REM transitions between other vigilance states. REM sleep had less theta power. Mice with synaptically blocked MnPO/MPO NOS1 neurons were also warmer. In particular, mice were warmer than control mice at the dark-light transition (ZT0), as well as during the dark phase siesta (ZT16-20), where there is usually a body temperature dip. Also, at this siesta point of cooled body temperature, mice usually have more NREM, but mice with synaptically blocked MnPO/MPO NOS1 cells showed reduced NREM sleep at this time. Overall, MnPO/MPO NOS1 neurons promote both NREM and REM sleep and contribute to chronically lowering body temperature, particularly at transitions where the mice normally enter NREM sleep.

## INTRODUCTION

Numerous circuits dispersed throughout the brain induce NREM sleep, but the preoptic (PO) hypothalamus, one of the first sleep-promoting centres to be identified (Nauta, 1946), has a major role (Sherin et al., 1996; Zhang et al., 2015; Weber and Dan, 2016; Chung et al., 2017; Kroeger et al., 2018; Ma et al., 2019; Reichert et al., 2019; Reitz and Kelz, 2021). The PO area also contains neurons that are required for REM sleep (Lu et al., 2000), including REM-promoting cells in MPO (Suntsova and Dergacheva, 2004; Gvilia et al., 2006). The PO area, which contains a huge diversity of cells (Moffitt et al., 2018; Tsuneoka and Funato, 2021), also contributes to regulating many other functions, including nesting, thermoregulation, parenting, sexual behaviour, water consumption, blood osmolarity and daily torpor (Nakamura and Morrison, 2008; Nakamura and Morrison, 2010; Morrison and Nakamura, 2011; Saper and Lowell, 2014; Abbott and Saper, 2017; Hrvatin et al., 2020; Takahashi et al., 2020; Tsuneoka and Funato, 2021).

The medial (M) and median (Mn) PO hypothalamic areas are enriched for neuronal nitric oxide (*Nos1*) gene expression, as seen by *in situ* hybridisation in the Allen Brain Atlas (Lein et al., 2007), and from our previous studies (Harding et al., 2018). Previously we found that *Nos1* neurons in the MnPO and MPO area link NREM onset and the decrease of body temperature that accompanies sleep (Harding et al., 2018). We hypothesised that external warm sensing and NREM sleep induction through these neurons may be part of an energy conservation mechanism that optimises sleep towards thermoneutral temperatures (Harding et al., 2020).

In addition to presumably synthesizing NO in response to excitation and calcium, MnPO/MPO NOS1 cells, depending on subtype, likely release both GABA and glutamate and/or various peptides (Moffitt et al., 2018). Here we show by calcium photometry that NOS1 neurons in MnPO/MPO have their highest activity during NREM sleep. Synaptic silencing of MnPO/MPO NOS1 cells with tetanus toxin light-chain (TeLC) expression induced bidirectional changes to NREM sleep structure: over the 24-hour cycle, mice had less NREM sleep in the dark phase, and more in the light phase. Dark phase REM sleep also consolidated to longer episodes, with a reduction in REM transitions; however, both light- and dark-phase REM sleep had more delta and less theta power than in controls, possibly suggesting disrupted REM function. In addition, a shift in the core body-temperature profile to warmer temperatures and a disrupted siesta (ZT16-20) period were observed. Thus, vesicular release of transmitters from MnPO/MPO hypothalamic NOS1 neurons is needed for maintaining normal sleep and temperature profiles.

## MATERIALS AND METHODS

### Mice

Experiments were performed under the Animals (Scientific Procedures) Act (1986) and approved by the local ethics committee. The mice used were *Nos1-ires-Cre^tm1(cr**E**)Mgmj^/J* (JAX labs stock 017526), referred to here as *Nos1-Cre* mice, donated by Martin G Myers (Leshan et al., 2012), and C57BL/6J mice (supplied by Charles River UK). All mice used in the experiments were male and congenic on a C57BL/6J background. Mice were maintained on a reversed 12 hr:12 hr light:dark cycle at constant temperature and humidity with *ad libitum* food and water.

### AAV transgenes

AAV transgenes were packaged in house into capsids with a 1:1 ratio of AAV1 and AAV2 capsid proteins. The adenovirus helper plasmid *pFΔ6*, the AAV helper plasmids *pH21* (AAV1) and *pRVI* (AAV2), and the *pAAV* transgene plasmids were co-transfected into HEK293 cells and AAVs harvested on heparin columns, as described previously (Klugmann et al., 2005; Yu et al., 2015). We used the following *pAAV* transgene plasmids: *pAAV-FLEX-GFP-TeLC* (Murray et al., 2011), and *pAAV-FLEX-GFP* (Addgene #28304, a gift from Edward Boyden). Plasmid *pAAV-FLEX-GCaMP6s* was created by inserting the *GCaMP6s* open reading frame from *pCMV-GCaMP6s* (Addgene plasmid 40753, gift of Douglas Kim, (Chen et al., 2013) into the backbone of *pAAV-flex-hM3Dq-mCHERRY* (Krashes et al., 2011) in place of the *hM3Dq* sequence, but retaining the loxP sites.

### Surgeries and stereotaxic injections of AAV

Mice underwent their first surgery at 10-weeks old. The mice required two rounds of surgery including implantation of an abdominal temperature logger, followed one week later by stereotaxic injections of AAV virus and electrode placement for electrocorticography (ECoG). For surgery, mice were anesthetized with 2% isoflurane and given appropriate analgesia. Viral infusions were performed using a steel injector (10 μl-Hamilton #701) and the aid of an electronic pump. Injections were optimized for the target with injection volumes of between 0.05 μl to 0.2 μl at 0.1 μl min^−1^. The injection coordinates relative to Bregma were AP +0.34 mm, ML 0 mm, DV −4.8 & 5.2. A minimum of one week recovery was allowed before recording the EEG.

### EEG and EMG recordings, scoring of vigilance states and power spectrum analysis

EEG and EMG were recorded from non-tethered animals using Neurologger 2A devices as described previously and electrodes placed at the same positions as our previous work in mice (Anisimov et al., 2014; Zhang et al., 2015). These positions were: AP +1.5 mm, ML −1.5 mm relative to Bregma, 1st - AP −1.5 mm, ML +1.5 relative to Bregma, 2nd Lambda −1.0 mm, ML 0.0 mm. EMG wires were also implanted in the neck muscles. Data were recorded at a sampling rate of 200 Hz with four times oversampling. The EEG data analyzed using Spike2 software 7.18 (Cambridge Electronic Design, Cambridge, UK) or MATLAB (MathWorks, Cambridge, UK). Prior to sleep scoring the ECoG was digitally filtered (high-pass, 0.5 Hz, −3dB) and the EMG was band-pass filtered (5-45 Hz, −3dB). Power in the delta (1-4 Hz) and theta (6-9 Hz) bands was calculated, together with the RMS value of the EMG signal (averaged over 5 s), and these were used to define the vigilance states of wake, NREM and REM with an automatic script OSD7 v7.2 (in Spike2). Each vigilance state was then rechecked manually. We analyzed the sleep-state specific power spectrums following normalization to wake power within each mouse, as described previously (Ma et al., 2019).

### Photometry recordings

Photometry was performed using a 473-nm diode-pumped solid state (DPSS) laser with fiber coupler (Shanghai Laser & Optics century Co.) and adjustable power supply (Shanghai Laser & Optics century Co.), controlled by a Grass SD9 stimulator. A lock-in amplifier (SR810, Stanford Research Systems, California, USA) drove the laser using a TTL signal at 125 Hz with an average power of 80 μW at the tip of the fiber. Using an optical fiber patch cord (Ø 200 μm, 0.22 NA, Doric Lenses) the light source passed through a fluorescence cube (FMC_GFP_FC, Doric Lenses) and then via a second optical patch cord (Ø 200 μm, 0.37 NA, Doric Lenses), was connected to the brain-implanted fiber via a ceramic sleeves (Thorlabs). The GCaMP6s output was then filtered at 500-550 nm (using a fluorescence cube) and passed to a photodiode (APD-FC, Doric Lenses) and amplified by the lock-in amplifier (time constant, 30 ms). The signal was recorded on a CED 1401 Micro box (Cambridge Electronic Design, Cambridge, UK) at 200 Hz using Spike2 software (Cambridge Electronic Design, Cambridge, UK). Photometry, EEG and EMG data were aligned offline using Spike2 and analyzed using custom scripts in either MATLAB (MathWorks) or R scripts (R Core Team, 2020). For each experiment, the photometry signal *F* was normalized to baseline using the function *ΔF/F* = *(F-F_0_)*/*F_0_*, where *F_0_* was the mean fluorescence across the signal analyzed.

### Temperature recordings

Core body temperatures were measured using an abdominally implanted temperature loggers (DSTnano, Star-Oddi, Herfølge, Denmark), sampling every two minutes, as described previously (Harding et al., 2018).

### Immunohistochemistry

Mice were given pentobarbital (100 mg/kg body weight; *i.p*.), and transcardially perfused with 4% paraformaldehyde in phosphate-buffered saline (PBS), pH 7.4. Brains were removed and 40-μm-thick coronal sections cut using a Leica SM 2010R microtome. Staining was performed on free-floating sections, washed in PBS three times and permeabilized in PBS plus 0.4% Triton X-100 for 30 min, blocked by incubation in PBS plus 10% normal goat serum (NGS), 0.2% Triton X-100 for 1 h (all at room temperature) and subsequently incubated overnight with a 1:1000 dilution of anti-GFP polyclonal antibody (A-6455, ThermoFisher). Sections were washed three times in PBS before incubating with goat anti-Rabbit IgG (H+L) Secondary, Alexa Fluor® 488 conjugate (A-11034, ThermoFisher) for two hours. Samples were then washed six times before mounting on VECTASHIELD Antifade Mounting Medium with DAPI (H-1200, Vector Laboratories).

### Statistics

Data collection and drug dosing were either randomized or performed in a counter-balanced manner. Data are represented as the mean ± SEM, unless otherwise stated. OriginPro 2017 was used for statistical analyses. For data that were not independent (where ANOVA was not appropriate) we employed either two-tailed or paired t-tests and then corrected for multiple comparisons using the Benjamini-Hochberg procedure at a false discovery rate of 5%. Mice were excluded from the analysis if the histology did not confirm AAV transgene expression in the MnPO/MPO area, or if the expression had spread beyond the target region. Investigators were not blinded to behavioral treatment groups.

## RESULTS

### Medial preoptic NOS1 neurons are most active during NREM sleep

We used calcium photometry to assess the sleep-wake activity of NOS1 neurons in the MPO area. *AAV-FLEX-GCaMP6s* was injected into the MnPO/MPO area of *Nos1-Cre* mice to generate *Nos1-MnPO/MPO-GCaMP6s* mice (**Figure 1A & 1B**). We then recorded calcium photometry signals from mice over six hours while the mice behaved freely in their home cages. Many NOS1 neurons in the MnPO/MPO region were NREM sleep-active, having their highest calcium activity in NREM sleep, and the cells were silent during wakefulness. An example over a six-minute period of a transition to NREM sleep is shown in **Figure 1C**, alongside the raw photometry signal, delta power (1-4 Hz), spectrogram from 0-20 Hz, EEG, EMG and scored sleep state. During wakefulness only low-level calcium-induced fluorescence signal was seen (labelled “F” on the axis of **Figure 1C**), and peaks in the signal were rare. While occasional small peaks in the calcium signal occurred during wake, a specific increase in peak frequency in the calcium signal was associated with NREM sleep. This is shown as a raster plot for ten transitions in **Figure 1D** over a six-minute period and quantified in **Figure 1E**. Higher GCaMP6s signal levels and more frequent peaks occurred during NREM sleep. Four example photometry traces are shown in **Figure 1F** and color coded by sleep state. Peak counting is shown above each example.

**Figure 1.**
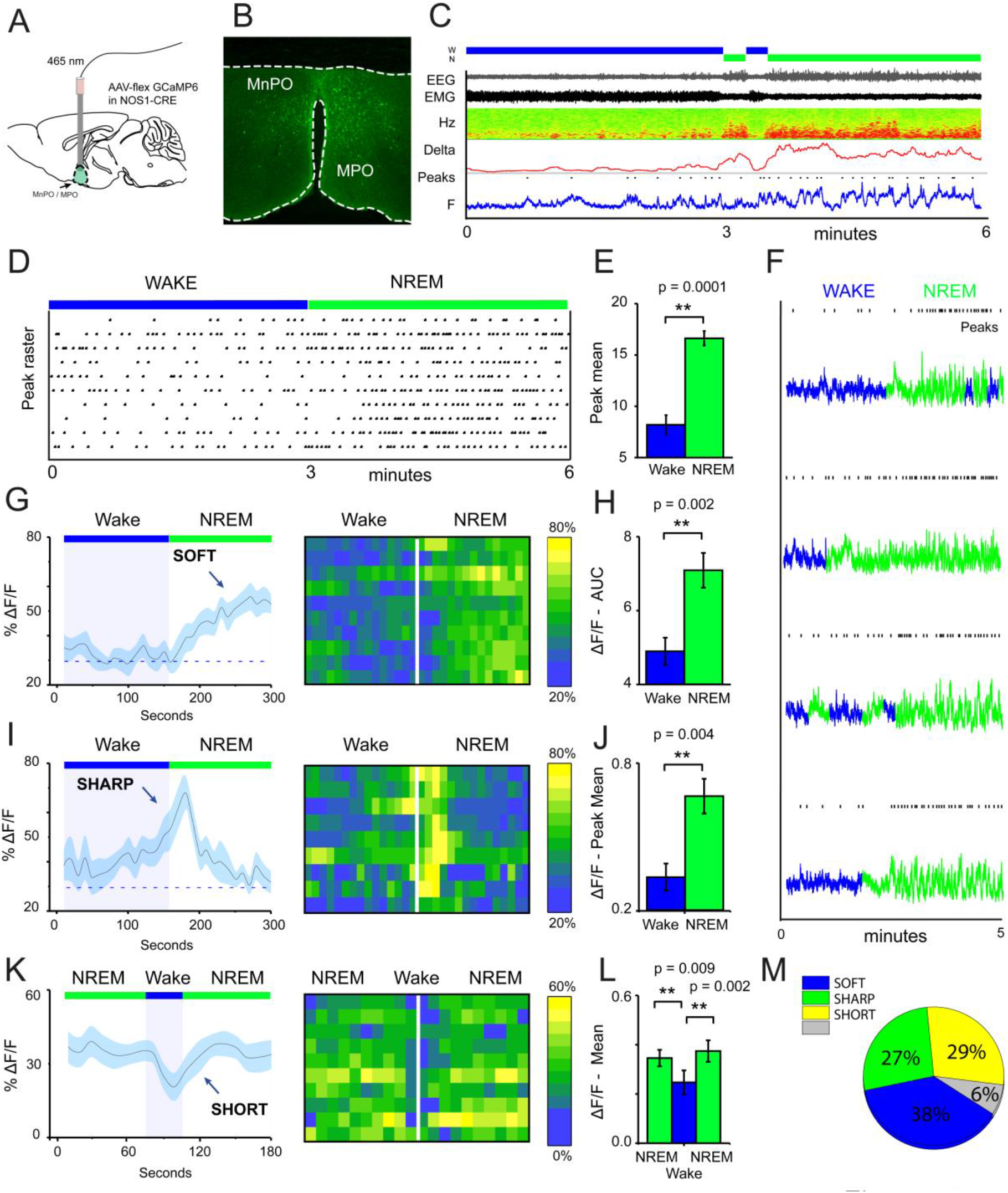
MnPO/MPO hypothalamic NOS1 neurons are more active during sleep. Animals were recorded for six hours across the light cycle from lights off to lights on to facilitate a distribution of sleep states. **(A)** Schematic for the photometry recording at a 5-mm depth and example of the GCaMP6 expression site in a *Nos1-MnPO/MPO-GCaMP6s* mouse. **(B)** Expression of GCaMP6s in neurons in the MnPO/MPO hypothalamus as detected by immunocytochemistry with GFP antisera. **(C)** Example transition from wake to NREM over a six-minute interval. Also shown are scored sleep states (Wake; W, NREM; N, REM; R), Filtered EEG and EMG, spectrogram of power in the frequency domain over time (Hz), Delta power (1-4 Hz) with a 5-second root mean square (RMS), raw photometry signal (labelled F) and automated peak counting on the photometry signal (Peaks). **(D)** A raster plot of automated spike counting from photometry signals across wake to NREM transitions. **(E)** area under the curve (ΔF/F) between wake and NREM for soft transitions (two-tailed t-test, n = 6, p = 0.0001). **(F)**Raw photometry data with paired automated peak counting for four example transitions over five minutes, colored by sleep state. Wake is shown in blue and NREM is shown in green. Peaks are marked above each trace. **(G)** The average ΔF/F for soft-type transitions that increase across wake-NREM transitions and ten example transitions represented as a heatmap. **(H)** ΔF/F Area under the curve between wake and NREM for soft transitions (two-tailed t-test, n = 6, p = 0.002). **(I)** The average ΔF/F for sharp-type transitions that increase in wake-NREM transitions before returning to baseline, followed by ten example transitions represented as a heatmap. **(J)** Mean change in ΔF/F between wake and NREM for sharp transitions (two-tailed t-test, n = 6, p = 0.004). **(K)** The average ΔF/F for SHORT-type transitions where high ΔF/F during NREM is interrupted by short bouts of WAKE but cannot return to baseline before the proceeding NREM episode. Wake length here is approximated and followed by ten example transitions represented as a heatmap. **(L)** Mean change in ΔF/F between brief wakefulness, prior-NREM and post-NREM for SHORT transitions (two-tailed t-test, n = 6, prior-NREM vs brief wakefulness, p = 0.009, post-NREM vs brief wakefulness p = 0.002). **(M)** Proportion of each transition type found in all transitions. Transitions that could not be classified are labelled NA (6%). All statistical tests were corrected for multiple comparisons using the Benjamini-Hochberg procedure at a false discovery rate of 5%.

On transitioning to NREM sleep the overall calcium levels in MnPO/MPO NOS1 neurons increased, as did the frequency of peaks. To quantify these changes in calcium signals, we averaged across multiple wake-NREM transitions from multiple mice. These averages contained several profiles: a slower ‘soft’ rising transition in calcium signal with more peaks; and a faster ‘sharp’ profile. Soft transitions from wake to NREM sleep are shown in **Figure 1G**, plotted as ΔF/F and averaged across five minutes of recording, alongside a heat map of ten example transitions. The ΔF/F calcium signal started to rise from the point of transition until it reached a plateau within about 100 seconds. This is quantified as the area under the curve in **Figure 1H**. On the other hand, sharp transitions in calcium-induced signals were different and anticipated the next NREM sleep transition. These sharp transitions are shown in **Figure 1I** as ΔF/F, alongside a heatmap of ten example transitions. Here, the ΔF/F signal started to rise up to 60 seconds prior to the start transition and peaked within 30 seconds of entering NREM sleep, before reducing again by 1 min post-transition. The heatmap shows that these events were time-locked to the wake-NREM transitions. Within 100 seconds, the calcium signal had returned to baseline, despite continuous NREM. This was quantified as the mean peak value in **Figure 1J**. In addition to the soft and sharp transitions in calcium signals of MnPO/MPO NOS1 neurons, NREM sleep was sometimes interrupted with short bouts of wake episodes lasting about one minute. In these short transitions, a small decrease in the calcium signal was seen on entry into wakefulness that continued to decline until the next NREM episode occurred and the signal increased once more (**Figure 1K**). This is shown alongside a heatmap of ten examples and quantified in **Figure 1L**. In this case, the signal during the short wake episodes was different to both adjacent NREM episodes. Both the soft and sharp transitions had a slow time course that took more than 60 seconds to complete. Overall, approximately 40% of transitions were soft and a further 30% were sharp. The remainder of transitions consisted of short bouts of wake that interrupted longer episodes of NREM. Approximately 5% of transitions did not have a clear association with the calcium signal (**Figure 1M**).

### Medial preoptic NOS1 neurons influence sleep-wake structure

Having established that many NOS1 neurons in the MnPO/MPO area are more active during NREM sleep, we next examined their contribution to sleep structure. To do this we reduced synaptic transmission from these cells, using Cre-dependent expression of tetanus-toxin light-chain (GFP-TeLC) (**Figure 2A**). Tetanus-toxin light chain blocks release of neurotransmitter vesicles by cleaving synaptobrevin, a synaptic vesicle protein (Schiavo et al., 1992). *AAV-FLEX-GFP-TeLC* or *AAV-FLEX-GFP* were injected into the MnPO/MPO of *Nos1-Cre* mice to generate *Nos1-MnPO/MPO-GFP-TeLC* and control *Nos1-MnPO/MPO-GFP* mice respectively (**Figure 2B**). Reducing synaptic transmission from NOS1 neurons produced small but significant alterations to the structure of sleep (**Figure 2C**). These data are quantified for each mouse. Average wakefulness was reduced by almost 25% during the light phase, with a corresponding increase in NREM of approximately 10%. This was followed by an approx. 15 to 20% decrease in NREM during lights OFF. No clear changes were seen in REM sleep time.

**Figure 2.**
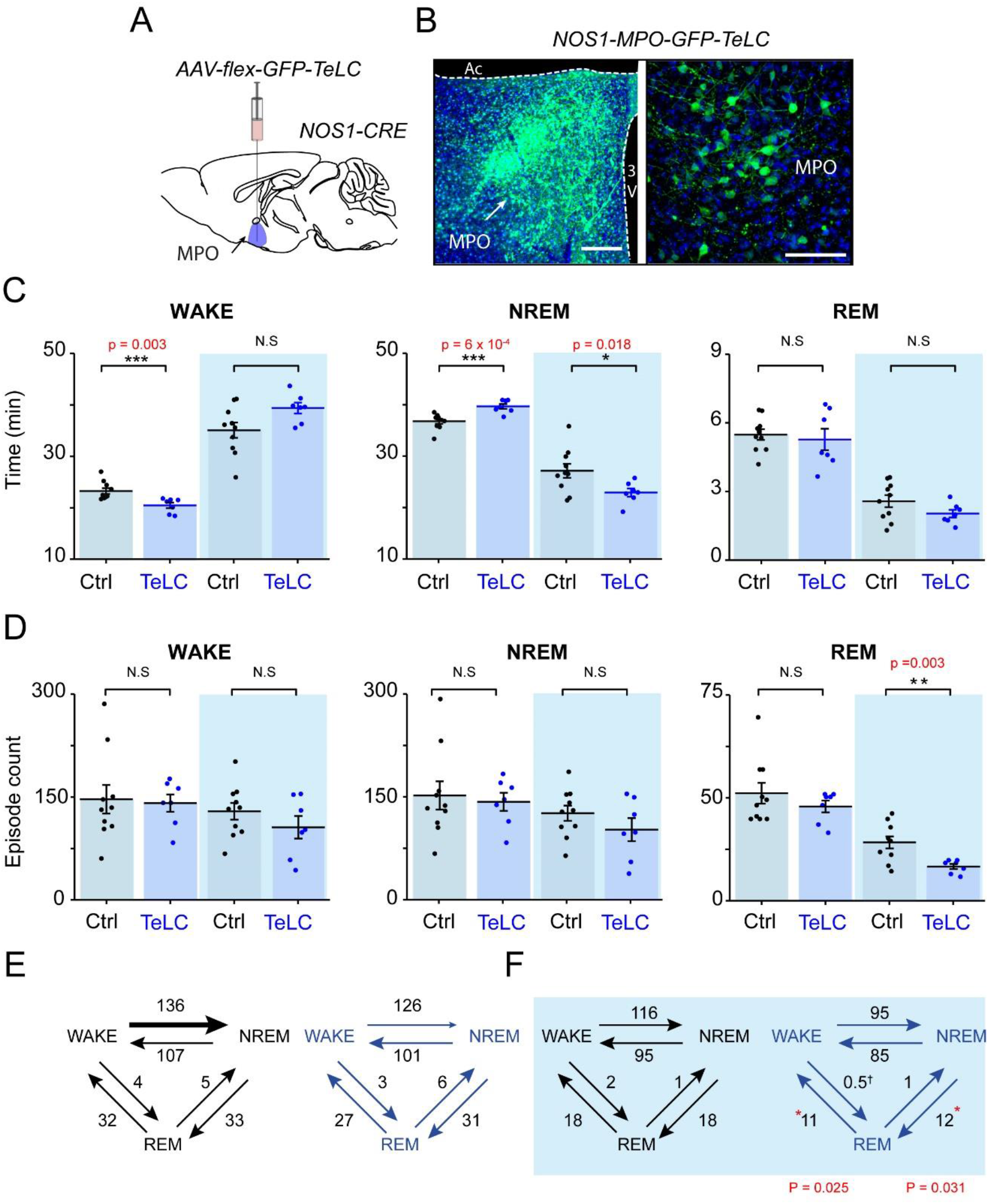
Reducing transmitter release from NOS1 neurons in MnPO/MPO hypothalamus alters sleep amounts and the number of sleep episodes in a manner dependent on the light-dark cycle. **(A)** Schematic of the stereotaxic injection of *AAV-flex-GFP-TeLC* into the MnPO/MPO area of *Nos1-CRE* mice to generate *Nos1-MnPO/MPO-GFP-TeLC* mice. **(B)** Example histology from MnPO/MPO showing expression of the GFP-TeLC protein as detected with a GFP antibody; left picture, lower magnification view, scale bar is 200 μm, right picture, higher magnification view, scale bar 100 μm **(C)** Quantification of sleep states for each mouse in the 12-hour light or dark periods shown as average time in vigilance state per hour. Wakefulness in the light phase (two-tailed t-test, n=7 and n=10, p = 0.003), NREM in the light phase (two-tailed t-test, n=7 and n=10, p = 0.0006), NREM in the dark phase (two-tailed t-test, n=7 and n=10, p = 0.018). **(D)** the number of episodes of wake, NREM and REM between light and dark. REM in the dark phase (two-tailed t-test, n=7 and n=10, p = 0.003). **(E)** Analysis of sleep transitions between sleep states in the light phase. No differences were observed between groups. **(F)** the transitions between *Nos1-MnPO/MPO-GFP-TeLC* mice and *Nos1-MnPO/MPO-GFP* mice in the dark phase. Transitions from NREM to REM (p = 0.025) and from REM to wake, p = 0.031), from two-tailed t-test, n=7 and n=10). All statistical tests were corrected for multiple comparisons using the Benjamini-Hochberg procedure at a false discovery rate of 5%.

We assessed whether changes in sleep structure seen in *Nos1-MnPO/MPO-GFP-TeLC* mice affected sleep episode dynamics and/or transitions (**Figure 2D**). There were no changes in the overall number of episodes in wake or NREM sleep for either the light or dark phase of the cycle; however, there was an approximately 45% reduction in REM episodes in the dark phase. **(Figure 2D, 2E)**. Consistent with this result, the number of NREM-REM and REM-wake transitions, but not transitions between wake and NREM sleep, were reduced by approximately 40%. Although this was not reflected in the REM sleep amount, it was consistent with less NREM in the dark phase. In addition, we expected the remaining REM sleep to be consequently more consolidated.

Because we did not observe an overall change in the number of wake or NREM episodes during the light or dark phase, we looked at the episode length and number of vigilance states to see if this explained the differences seen in the time spent sleeping (**Figure 3**). In *Nos1-MnPO/MPO-GFP-TeLC* mice we observed reductions of approximately 50% in the frequency of the longest wake episodes (>20 min) in the light phase (**Figure 3A**), although no changes were seen in episode length and number for NREM and REM sleep (**Figure 3B and C**). In the dark phase, the average values for wake did not change, and the data had larger variance (**Figure 3D**). However, while *Nos1-MnPO/MPO-GFP* control mice had a reduced frequency of episodes, > 20min in the dark phase compared with the light phase, the *Nos1-MnPO/MPO-GFP-TeLC* mice instead had an increased frequency of these episodes. From calculating the paired difference for each mouse between the light and dark phase, it was clear that *Nos1-MnPO/MPO-GFP-TeLC* mice were more affected by the light change (**Figure 3D**, inset graph). No further alterations were seen in episode length and number for NREM and REM sleep during lights OFF (**Figure 3E and F)**.

**Figure 3.**
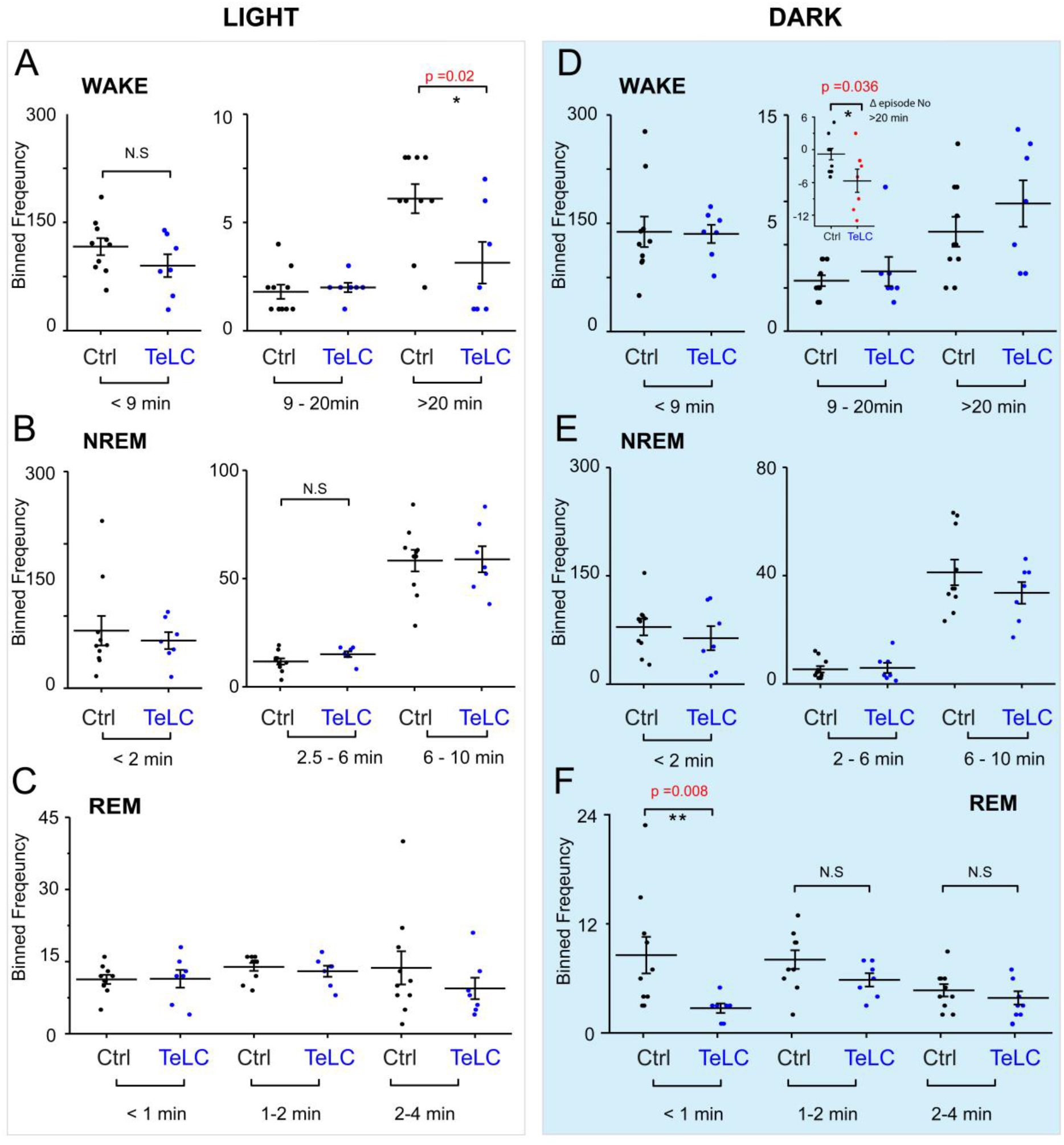
Distribution of sleep episode lengths is reduced for *Nos1-MnPO/MPO-GFP-TeLC* mice during wakefulness in the light period but increased in wakefulness in the dark period. (**A**) Wakefulness in the light period binned by episode length. No differences are seen in bins less than 20 mins. Episodes greater than 20 mins in *Nos1*-*MnPO/MPO-GFP-TeLC* mice (n = 7 TeLC and n =10 GFP, p = 0.02). **(B)** NREM in the light period binned by episode length. No differences are seen in bins less than 10 min (n = 7 TeLC and n =10 GFP). **(C)** light phase REM. No differences are seen in bins less than 4 mins (n = 7 TeLC and n =10 GFP). **(D)** dark phase wakefulness. No differences are seen between groups within each light cycle (n = 7 TeLC and n =10 GFP). Inset graph, between the light periods *Nos1*-*MnPO/MPO-GFP* and *Nos1*-*MnPO/MPO-GFP-TeLC* groups as a paired difference (d, inset graph, n = 7 TeLC and n =10 GFP, p = 0.036). **(E)** dark phase NREM. No differences are seen in bins less than 10 mins (n = 7 TeLC and n =10 GFP). **(F)** dark phase REM. No differences are seen in bins greater than 1 min. Episodes of less than 1 min are significant between *Nos1-MnPO/MPO-GFP-TeLC* and *Nos1-MnPO/MPO-GFP* mice (n = 7 TeLC and n =10 GFP, p = 0.008).

To better illustrate the differences in sleep structure, example data from a *Nos1-MnPO/MPO-GFP-TeLC* and *Nos1-MnPO/MPO-GFP* mouse are shown in **Figure 4**, plotted as episode length (log scale) vs the episode sequence number (the position the episode occurred in the sleep recording irrespective of length). The light-cycle example for wakefulness is shown in **Figure 4A**. The dotted line at 20 mins indicates episode length bins associated with a decreased number of longer-episodes in the *Nos1-MnPO/MPO-GFP-TeLC* mouse compared with *Nos1-MnPO/MPO-GFP (see also Figure 3A)*. Wake-sleep during the dark phase is shown in **Figure 4B**, but we did not observe further changes in sleep. This was also true for NREM in both the light and dark phase shown in **Figure 4C** and **D**. While no changes were seen in REM structure during the light phase (**Figure 4E**), large reductions in the number of short episodes of REM sleep in the dark phase were seen (**Figure 4F**). The points for comparison are highlighted by the dotted lines.

**Figure 4.**
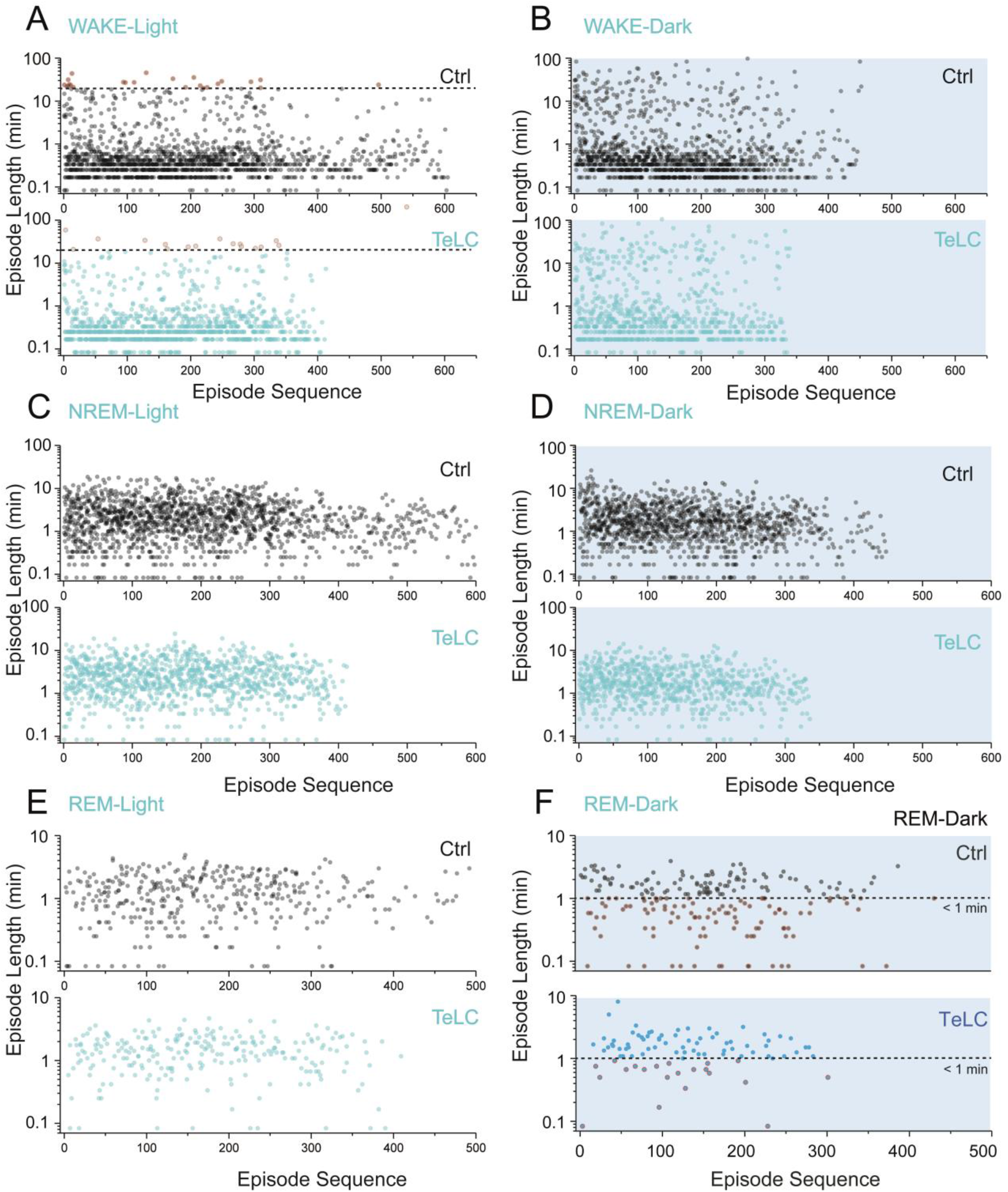
Example comparison of raw data from an individual *Nos1-MnPO/MPO-GFP-TeLC* and *Nos1-MnPO/MPO-GFP* mouse highlighting the decreased long wake episodes and decreased short REM episodes that appear when vesicular transmitter release is blocked from MnPO/MPO NOS1 cells. **(A)** Wake during the light period. Points with red boundaries are different between *Nos1*-*MnPO/MPO-GFP* and TeLC mice. **(B)** Wake during the dark period. **(C)** NREM during the light period **(D)** NREM during Lights-OFF. **(E)** REM during the light period. **(F)** REM during the dark period. Points with red boundaries are different between *Nos1-MnPO/MPO-GFP* and *Nos1-MnPO/MPO-GFP-TeLC* mice. Raw data from an individual *Nos1-MnPO/MPO-GFP-TeLC* mouse compared with a control *Nos1*-*MnPO/MPO-GFP* mouse. Episode length is displayed in minutes on a log scale. Dashed lines represent y-axis boundaries for cases where differences were observed in **(A)** and **(F)**.

### Medial preoptic NOS1 neurons contribute to theta power during REM sleep

We analyzed the sleep-state specific power spectra of *Nos1-MnPO/MPO-GFP-TeLC* mice compared with *Nos1-MnPO/MPO-GFP* control mice, following normalization to WAKE power within each mouse. During the lights-on phase, NREM sleep was not associated with changes in power (**Figure 5A and B**); however, REM sleep did show significant changes (**Figure 5C**). Specifically, there was an increase in delta power, normally associated with NREM sleep, of approximately 30%, as well as a corresponding decrease in theta power (**Figure 5D**). This difference was in the 2-4 Hz range of delta power referred to as the δ2 band (Hubbard et al., 2020). Theta (6-9 Hz) power was reduced by approximately 20% but no changes in the higher frequencies (10-14 Hz) were seen. During the light phase (**Figure 5E and F**), NREM showed an approximately 15% reduction in theta power. This contrasted with no change in this band during the light phase. For REM sleep in the dark phase (**Figure 5G and H**), differences between *Nos1-MnPO/MPO-GFP-TeLC* and *Nos1-MnPO/MPO-GFP* mice mirrored those seen in the light phase, with an approximately 25% increase in δ2 power as well as a 20% decrease in theta power.

**Figure 5.**
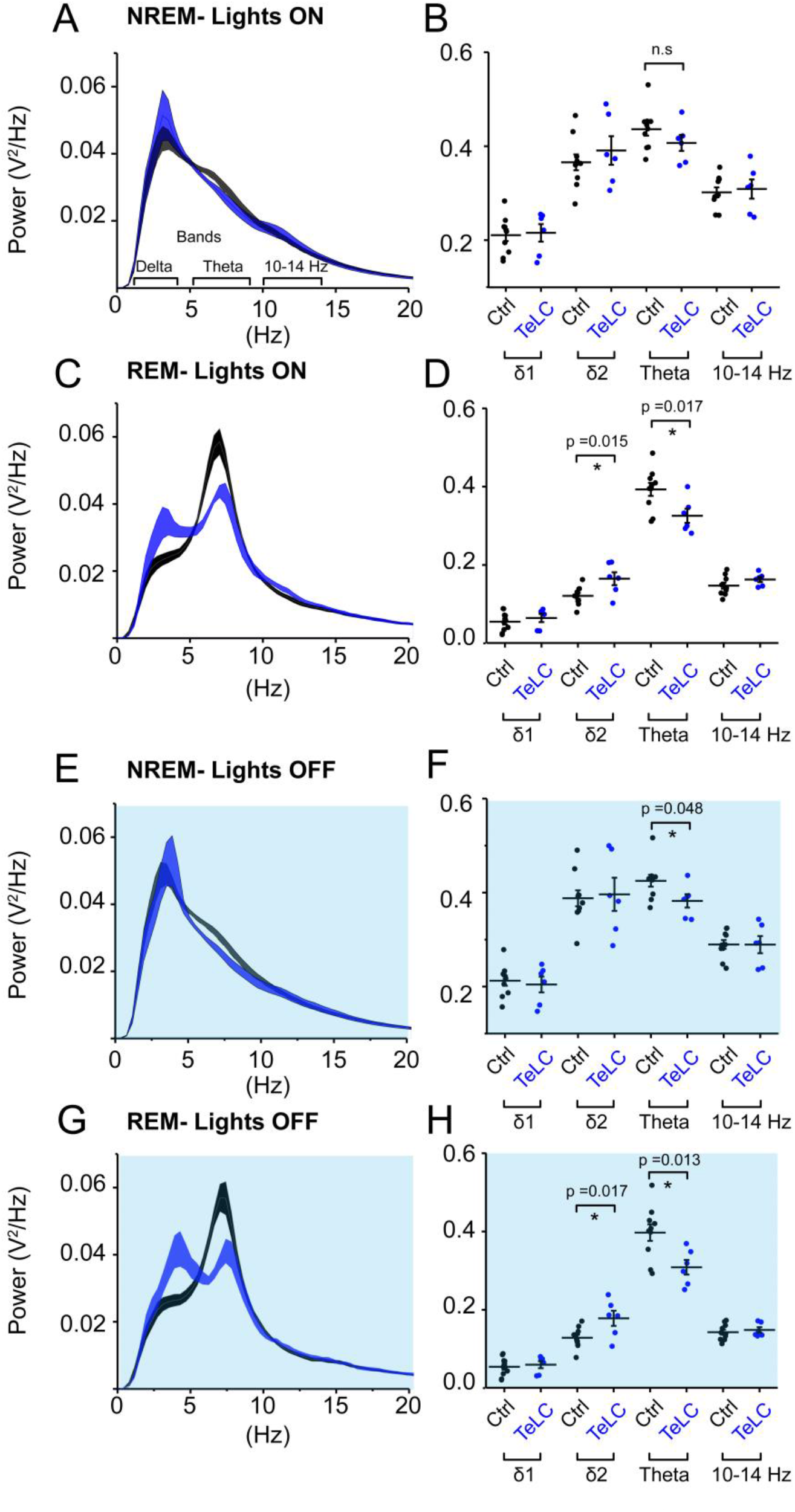
NOS1 neurons in MnPO/MPO contribute to theta power during REM sleep. **(A)** Normalized power for NREM during the light period shown as a power spectrum. Different frequency bands are also illustrated **(B)** For NREM during the light period, data for individual mice are shown for relevant frequency bands. **(C)** REM during the light period **(D)** Quantification of spectral differences during REM in the light period for individual mice show more δ2 in the TeLC condition (t test, n = 6, TeLC; n=10, GFP; p = 0.015) alongside reduced theta (t test, n = 6, TeLC; n=10, GFP; p = 0.017). **(E)** Normalized power for NREM in the dark period shown as a power spectrum. **(F)** Data for individual mice are shown for relevant frequency bands (t test, n = 6, TeLC; n=10, GFP; p = 0.048). **(G)** REM during the dark period **(H)** Quantification of spectral differences during REM in the light period for individual mice (δ2 between groups, t test, n = 6, TeLC; n=10, GFP; p = 0.017). Theta between groups (t test, n = 6, TeLC; n=10, GFP; p = 0.013). All statistical tests were corrected for multiple comparisons using the Benjamini-Hochberg procedure at a false discovery rate of 5%.

### Medial preoptic NOS1 neurons reduce body temperature

Using implanted temperature loggers, we measured the core body temperature at 2-min resolution in *Nos1-MnPO/MPO-GFP-TeLC* and *Nos1-MnPO/MPO-GFP* mice (**Figure 6**). We produced a typical 24-hour period in temperature change by first averaging over seven days for each mouse (5040 measurements) before comparing distributions across groups. The temperature distribution of *Nos1-MnPO/MPO-GFP-TeLC* mice shifted to warmer temperatures compared with those of control mice (**Figure 6A**). The cumulative distribution illustrated that the most significant change was in the probability of observing core temperature between 35.5 to 36°C, but without a change in the minimum or maximum temperatures (**Figure 6B**). Furthermore, during the dark phase, while *Nos1-MnPO/MPO-GFP* control mice dropped their core temperature, *Nos1-MnPO/MPO-GFP-TeLC* control mice did not, both during the middle of the dark phase and prior to the next lights-on period (**Figure 6C, D**); furthermore, during the siesta period *Nos1-MnPO/MPO-GFP-TeLC* mice spent 36% less total time in NREM sleep compared with control mice, although there were no significant changes in NREM sleep episode count or mean length (**Figure 6E**).

**Figure 6.**
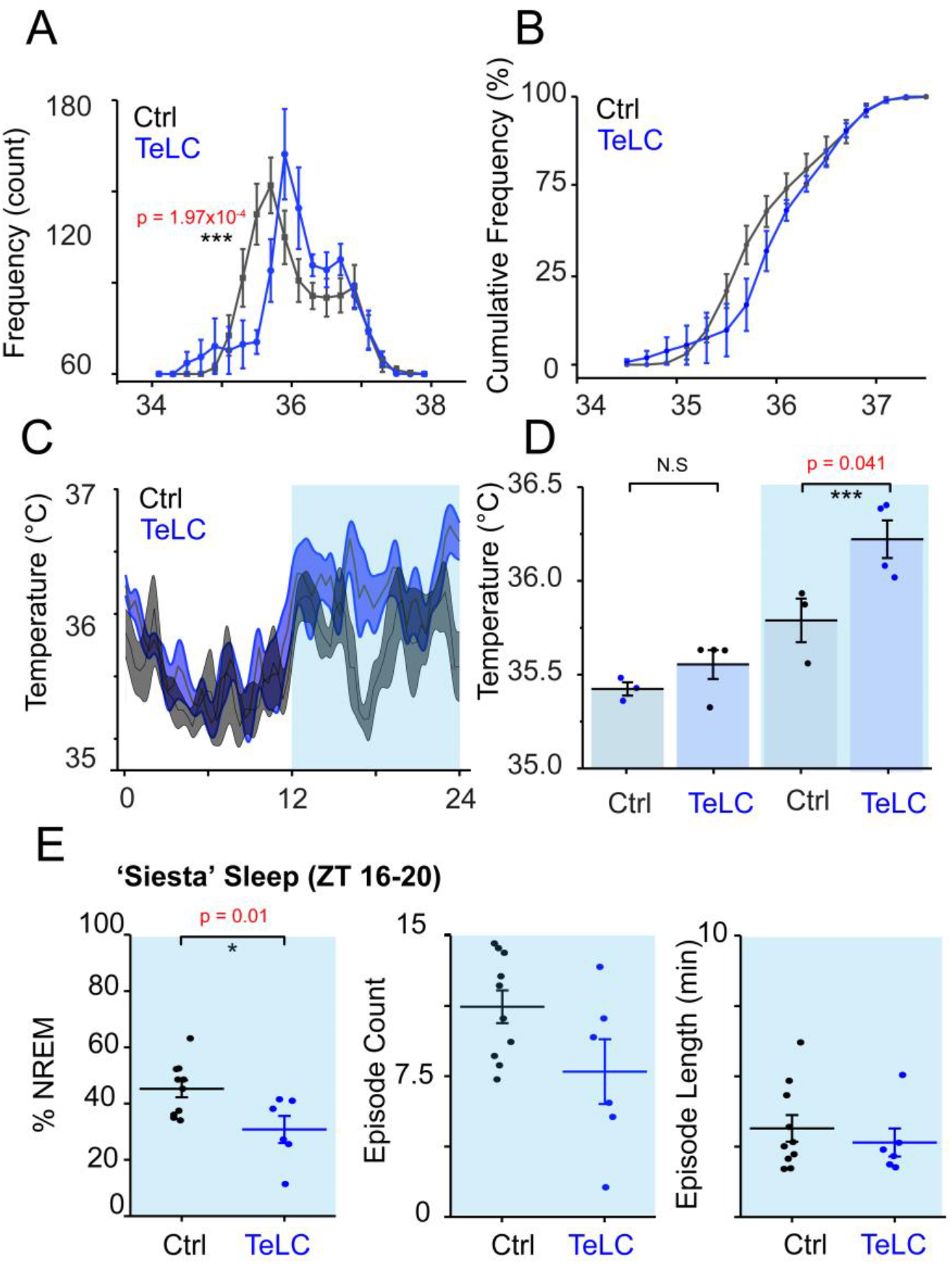
NOS1 neurons in MnPO/MPO hypothalamus act to reduce body temperature. **(A)** Aggregated temperature data from 7 days of recording for each mouse averaged by ZT time at 2-min intervals to produce a ‘typical’ day histogram, averaged for each group. (rmANOVA, n = 4, TeLC; n=6, GFP; temperature*group at 35.4-35.6°C, p=1.97 × 10^−4^). **(B)** Cumulative frequency of temperature distribution **(C)** Temperature profile over 24 hours. **(D)** Quantification of 24-hour temperature profile in the light and dark phase (two-tailed t test assuming unequal variance, n = 4, TeLC; n=3, GFP; p = 0.041). **(E)** Percentage sleep and episode count and episode length a ‘siesta’ period between ZT 16-20 (two tailed t-test, n=7 and n=10, p = 0.01).

## DISCUSSION

In this study we have discovered that many NOS1 cells in the midline PO hypothalamus are naturally sleep-active. To our knowledge this is the first instance of a specifically NREM-active population in the hypothalamus. In terms of their calcium signals, the fast and transient ‘sharp’ activations of these NOS1 cells from wake to NREM and the slower, prolonged ‘soft’ transitions from wake to NREM may represent two populations of sleep-active NOS1 neurons. We used TeLC expression in NOS1 neurons to disrupt their synaptic activity, which in turn disrupted the sleep-wake profile of the mice in a manner that varied with the light or dark phases of the 24-hour cycle. In the dark phase, mice with TeLC expressed in MnPO/MPO NOS1 neurons showed a reduction of time in NREM sleep and a loss of the shortest REM episodes; NREM to REM and REM to wake transitions were also reduced. REM sleep was accompanied by increased delta power and decreased theta power, possibly suggesting functional disruption of REM sleep. In the light phase, however, there was an increase in NREM sleep and REM sleep was unchanged. Overall, the mice were chronically warmer.

Our new results are consistent with our previous work on these cells. We have previously shown that a subset of MnPO/MPO glutamate/NOS1 neurons, when activity-tagged following an external warm-stimulus to the mice, could on reactivation induce NREM sleep and concomitant body cooling (Harding et al., 2018). Similarly, a GABAergic MnPO/MPO population, tagged in the same manner, could only induce sleep (Harding et al., 2018). As we did not observe overlap in these populations by immunohistochemistry, we suggested a model of external warmth-triggered sleep with a NOS1/glutamate (MnPO/MPO) population signaling to a downstream GABAergic population in MPO (Harding et al., 2020). The effects on REM sleep were unanticipated, but perhaps not surprising given that REM sleep has been found to be partly controlled by unknown cell types in the MPO area (Suntsova and Dergacheva, 2004; Gvilia et al., 2006), and we have presumably influenced a NOS1 cell subtype involved in REM production.

Initially, given that *nos1* gene expression in the MPO hypothalamus has a highly restricted expression pattern, as detected by both *in situ* hybridization and immunocytochemistry (see e.g. Fig. 4E in Harding et al., 2018), we anticipated that *nos1* would be a pragmatic and useful marker for functional manipulation of a unique subset of cells. Unfortunately, this has turned out not to be the case. While many of the NOS1 neurons in MPO studied by calcium photometry have clear sleep-active patterns, it has become apparent since we started our work that multiple subtypes of NOS1 neuron exist in the PO area, including NOS1/VGLUT2, NOS1/VGAT, NOS1/galanin neurons and others (Moffitt et al., 2018). The bidirectional changes in sleep when TeLC is expressed in MnPO/MPO NOS1 neurons thus likely reflect reduced synaptic transmitter release from multiple subtypes of NOS1 cell in MnPO/MPO. For example, activation of glutamate (VGLUT2) neurons in the PO area induces wakefulness (Vanini et al., 2020), so if this particular subset were to express the *nos1* gene, TeLC expression in them might reduce wakefulness; on the other hand, we have shown previously that *nos1*-expressing GABA cells induce NREM sleep (Harding et al., 2018); thus TeLC expression in NOS1 cells might promote wakefulness (Harding et al., 2018). Further progress to dissect this circuitry requires intersectional genetics. Nevertheless, it remains striking that the majority of MnPO/MPO NOS1 cells have most of their activity during NREM sleep. Targets for NOS1 neurons could include GABAergic and galaninergic neurons in the LPO area that are involved in NREM sleep induction and maintenance (Kroeger et al., 2018; Ma et al., 2019), as well as uncharacterized long-range targets.

Expressing TeLC in MnPO/MPO NOS1 neurons raised the average body temperature of the mice. This would be consistent with effects on temperature mediated by BDNF/PACAP or TRPM2 expressing neurons in the MPO area (Song et al., 2016; Tan et al., 2016; Harding et al., 2018); these neurons could co-express NOS1. There are also glutamatergic wake-promoting neurons in the PO that are associated with mild body cooling of approximately 1°C that could also have a role (Vanini et al., 2020), and which also could express the *nos1* gene. However, unlike the effects on temperature produced by BDNF/PACAP or TRPM2 cells, the increases we see appear to be associated with the light phase of the dark-light cycle, specifically in the siesta period (ZT16-20) and the period before the dark-to-light transition. Thus, MnPO/MPO NOS1 cells are driving down temperature at the same time as NREM sleep is initiated, consistent with our earlier work (Harding et al., 2018). Overall, this may support a larger hypothesis on optimization of sleep for energy reallocation (Harding et al., 2020).

In summary, we have found that the activity pattern of MnPO/MPO NOS1 cells is quite striking, being rather selective either at the boundary between wake to NREM transitions and being sustained during NREM sleep, and that synaptic transmission from PO NOS1 neurons likely contributes to NREM and REM sleep organization, as well as chronic body cooling. We write “likely” because we have not formally shown that TeLC expression did reduce transmitter release, and we have not identified post-synaptic targets of MnPO/MPO NOS1 cells. We currently think that MnPO/MPO NOS1 neurons probably have both short local outputs and long-range connections where transmitters could be released. A further caveat is that NO itself is likely to be part of the signaling system from these cells. We did not address this because NO release from cells is independent of vesicle release. But as NOS1 synthase is calcium-dependent (Knowles and Moncada, 1994), periods of elevated calcium seen in NOS1 neurons at the wake to NREM transitions and during NREM sleep will result in NO release from these cells, and NO could well be influencing sleep structure and temperature regulation. Given their rather precise calcium activity at the boundary of wake to NREM transitions and during NREM asleep, further dissection of MnPO/MPO NOS1 cells will likely reveal part of a regulatory circuit controlling sleep induction/maintenance and the simultaneous lowering of body temperature.

## AUTHOR CONTRIBUTIONS

W.W., N.P.F. and E.C.H. conceived the project. E.C.H. with input from W.W., N.P.F. designed the experiments. E.C.H., W.B., R.Z., X.Y., R.Y., L.L performed the experiments. R.Z. and L.L. were supervised by E.C.H. for this work. B.H. and T.C. provided technology for EEG recordings. A.L.V. provided the Neurologgers. E.C.H. performed the data analysis and produced the figures. N.P.F. and W.W. contributed to the data analysis and supervised the project. E.C.H with N.P.F. and W.W. wrote the paper.

## ACKNOWLEDGMENTS

We are grateful to Stephen Rothery and David Gaboriau for technical assistance with the light microscopy.

## REFERENCES

Abbott, S.B.G., and Saper, C.B. (2017). Median preoptic glutamatergic neurons promote thermoregulatory heat loss and water consumption in mice. The Journal of physiology 595(20), 6569–6583. doi: 10.1113/JP274667.

Anisimov, V.N., Herbst, J.A., Abramchuk, A.N., Latanov, A.V., Hahnloser, R.H., and Vyssotski, A.L. (2014). Reconstruction of vocal interactions in a group of small songbirds. Nat Methods 11(11), 1135–1137. doi: 10.1038/nmeth.3114.

Chen, T.-W., Wardill, T.J., Sun, Y., Pulver, S.R., Renninger, S.L., Baohan, A., et al. (2013). Ultrasensitive fluorescent proteins for imaging neuronal activity. Nature 499(7458), 295–300. doi: 10.1038/nature12354.

Chung, S., Weber, F., Zhong, P., Tan, C.L., Nguyen, T.N., Beier, K.T., et al. (2017). Identification of preoptic sleep neurons using retrograde labelling and gene profiling. Nature 545(7655), 477–481. doi: 10.1038/nature22350.

Gvilia, I., Turner, A., McGinty, D., and Szymusiak, R. (2006). Preoptic area neurons and the homeostatic regulation of rapid eye movement sleep. J Neurosci 26(11), 3037–3044. doi: 10.1523/JNEUROSCI.4827-05.2006.

Harding, E.C., Franks, N.P., and Wisden, W. (2020). Sleep and thermoregulation. Current Opinion in Physiology 15, 7–13. doi: https://doi.org/10.1016/j.cophys.2019.11.008.

Harding, E.C., Yu, X., Miao, A., Andrews, N., Ma, Y., Ye, Z., et al. (2018). A Neuronal Hub Binding Sleep Initiation and Body Cooling in Response to a Warm External Stimulus. Curr Biol 28(14), 2263–2273.e2264. doi: 10.1016/j.cub.2018.05.054.

Hrvatin, S., Sun, S., Wilcox, O.F., Yao, H., Lavin-Peter, A.J., Cicconet, M., et al. (2020). Neurons that regulate mouse torpor. Nature 583(7814), 115–121. doi: 10.1038/s41586-020-2387-5.

Hubbard, J., Gent, T.C., Hoekstra, M.M.B., Emmenegger, Y., Mongrain, V., Landolt, H.P., et al. (2020). Rapid fast-delta decay following prolonged wakefulness marks a phase of wake-inertia in NREM sleep. Nat Commun 11(1), 3130. doi: 10.1038/s41467-020-16915-0.

Klugmann, M., Symes, C.W., Leichtlein, C.B., Klaussner, B.K., Dunning, J., Fong, D., et al. (2005). AAV-mediated hippocampal expression of short and long Homer 1 proteins differentially affect cognition and seizure activity in adult rats. Mol Cell Neurosci 28(2), 347–360. doi: 10.1016/j.mcn.2004.10.002.

Knowles, R.G., and Moncada, S. (1994). Nitric oxide synthases in mammals. Biochem J 298 (Pt 2), 249–258. doi: 10.1042/bj2980249.

Krashes, M.J., Koda, S., Ye, C., Rogan, S.C., Adams, A.C., Cusher, D.S., et al. (2011). Rapid, reversible activation of AgRP neurons drives feeding behavior in mice. J Clin Invest 121(4), 1424–1428. doi: 10.1172/jci46229.

Kroeger, D., Absi, G., Gagliardi, C., Bandaru, S.S., Madara, J.C., Ferrari, L.L., et al. (2018). Galanin neurons in the ventrolateral preoptic area promote sleep and heat loss in mice. Nat Commun 9(1), 4129. doi: 10.1038/s41467-018-06590-7.

Lein, E.S., Hawrylycz, M.J., Ao, N., Ayres, M., Bensinger, A., Bernard, A., et al. (2007). Genome-wide atlas of gene expression in the adult mouse brain. Nature 445(7124), 168–176. doi: 10.1038/nature05453.

Leshan, R.L., Greenwald-Yarnell, M., Patterson, C.M., Gonzalez, I.E., and Myers, M.G., Jr. (2012). Leptin action through hypothalamic nitric oxide synthase-1-expressing neurons controls energy balance. Nat Med 18(5), 820–823. doi: 10.1038/nm.2724.

Lu, J., Greco, M.A., Shiromani, P., and Saper, C.B. (2000). Effect of lesions of the ventrolateral preoptic nucleus on NREM and REM sleep. J Neurosci 20(10), 3830–3842.

Ma, Y., Miracca, G., Yu, X., Harding, E.C., Miao, A., Yustos, R., et al. (2019). Galanin Neurons Unite Sleep Homeostasis and alpha2-Adrenergic Sedation. Curr Biol 29(19), 3315–3322 e3313. doi: 10.1016/j.cub.2019.07.087.

Moffitt, J.R., Bambah-Mukku, D., Eichhorn, S.W., Vaughn, E., Shekhar, K., Perez, J.D., et al. (2018). Molecular, spatial, and functional single-cell profiling of the hypothalamic preoptic region. Science 362(6416). doi: 10.1126/science.aau5324.

Morrison, S.F., and Nakamura, K. (2011). Central neural pathways for thermoregulation. Frontiers in bioscience (Landmark edition) 16, 74–104. doi: 10.2741/3677.

Murray, A.J., Sauer, J.F., Riedel, G., McClure, C., Ansel, L., Cheyne, L., et al. (2011). Parvalbumin-positive CA1 interneurons are required for spatial working but not for reference memory. Nat Neurosci 14(3), 297–299. doi: 10.1038/nn.2751.

Nakamura, K., and Morrison, S.F. (2008). A thermosensory pathway that controls body temperature. Nat Neurosci 11(1), 62–71. doi: 10.1038/nn2027.

Nakamura, K., and Morrison, S.F. (2010). A thermosensory pathway mediating heat-defense responses. Proceedings of the National Academy of Sciences 107(19), 8848. doi: 10.1073/pnas.0913358107.

Nauta, W.J. (1946). Hypothalamic regulation of sleep in rats; an experimental study. J Neurophysiol 9, 285–316. doi: 10.1152/jn.1946.9.4.285.

R Core Team (2020). R: A language and environment for statistical computing. R Foundation for Statistical Computing, Vienna, Austria.

Reichert, S., Pavon Arocas, O., and Rihel, J. (2019). The Neuropeptide Galanin Is Required for Homeostatic Rebound Sleep following Increased Neuronal Activity. Neuron 104(2), 370–384 e375. doi: 10.1016/j.neuron.2019.08.010.

Reitz, S.L., and Kelz, M.B. (2021). Preoptic Area Modulation of Arousal in Natural and Drug Induced Unconscious States. Front Neurosci 15, 644330. doi: 10.3389/fnins.2021.644330.

Saper, C.B., and Lowell, B.B. (2014). The hypothalamus. Current Biology 24(23), R1111–R1116. doi: https://doi.org/10.1016/j.cub.2014.10.023.

Schiavo, G., Benfenati, F., Poulain, B., Rossetto, O., Polverino de Laureto, P., DasGupta, B.R., et al. (1992). Tetanus and botulinum-B neurotoxins block neurotransmitter release by proteolytic cleavage of synaptobrevin. Nature 359(6398), 832–835. doi: 10.1038/359832a0.

Sherin, J.E., Shiromani, P.J., McCarley, R.W., and Saper, C.B. (1996). Activation of ventrolateral preoptic neurons during sleep. Science 271(5246), 216–219. doi: 10.1126/science.271.5246.216.

Song, K., Wang, H., Kamm, G.B., Pohle, J., Reis, F.C., Heppenstall, P., et al. (2016). The TRPM2 channel is a hypothalamic heat sensor that limits fever and can drive hypothermia. Science 353(6306), 1393–1398. doi: 10.1126/science.aaf7537.

Suntsova, N.V., and Dergacheva, O.Y. (2004). The role of the medial preoptic area of the hypothalamus in organizing the paradoxical phase of sleep. Neurosci Behav Physiol 34(1), 29–35. doi: 10.1023/b:neab.0000003243.95706.de.

Takahashi, T.M., Sunagawa, G.A., Soya, S., Abe, M., Sakurai, K., Ishikawa, K., et al. (2020). A discrete neuronal circuit induces a hibernation-like state in rodents. Nature 583(7814), 109–114. doi: 10.1038/s41586-020-2163-6.

Tan, C.L., Cooke, E.K., Leib, D.E., Lin, Y.C., Daly, G.E., Zimmerman, C.A., et al. (2016). Warm-Sensitive Neurons that Control Body Temperature. Cell 167(1), 47–59.e15. doi: 10.1016/j.cell.2016.08.028.

Tsuneoka, Y., and Funato, H. (2021). Cellular composition of the preoptic area regulating sleep, parental, and sexual behavior *Frontiers in Neuroscience, section Sleep and Circadian Rhythms*.

Vanini, G., Bassana, M., Mast, M., Mondino, A., Cerda, I., Phyle, M., et al. (2020). Activation of Preoptic GABAergic or Glutamatergic Neurons Modulates Sleep-Wake Architecture, but Not Anesthetic State Transitions. Current Biology 30(5), 779–787.e774. doi: 10.1016/j.cub.2019.12.063.

Weber, F., and Dan, Y. (2016). Circuit-based interrogation of sleep control. Nature 538(7623), 51–59. doi: 10.1038/nature19773.

Yu, X., Ye, Z., Houston, C.M., Zecharia, A.Y., Ma, Y., Zhang, Z., et al. (2015). Wakefulness Is Governed by GABA and Histamine Cotransmission. Neuron 87(1), 164–178. doi: 10.1016/j.neuron.2015.06.003.

Zhang, Z., Ferretti, V., Guntan, I., Moro, A., Steinberg, E.A., Ye, Z., et al. (2015). Neuronal ensembles sufficient for recovery sleep and the sedative actions of alpha2 adrenergic agonists. Nat Neurosci 18(4), 553–561. doi: 10.1038/nn.3957.

